# Limitations of qPCR to estimate DNA quantity: An RFU method to facilitate inter-laboratory comparisons for activity level, and general applicability

**DOI:** 10.1101/2022.05.23.493102

**Authors:** Peter Gill, Øyvind Bleka, Ane Elida Fonneløp

**Affiliations:** Forensic Genetics Research Group, Oslo University Hospital, Oslo, Norway; Department of Forensic Medicine, University of Oslo, Oslo, Norway

**Keywords:** Average peak height, quantification, qPCR, Activity level, Normalisation

## Abstract

The application of qPCR to estimate the quantity of DNA present is usually based upon a short amplicon (typically c.80bp) and a longer amplicon (typically c.200-300bp) where the latter is used to determine the amount of degradation present in a sample. The data are used to make decisions about a) whether there is sufficient template to amplify? b) how much of the elution volume to forward to PCR? A typical multiplex amplifies template in the region of 100-500bp. Consequently, the results from an 80bp amplicon will tend to overestimate the actual amplifiable quantity that is present in a degraded sample. To compensate, a method is presented that relates the quantity of amplifiable DNA to the average RFU of the amplified fragments. This provides greatly improved accuracy of the estimated quantity of DNA present, which may differ by more than an order of magnitude compared to qPCR. The relative DNA quantities can be apportioned per contributor once mixture proportions are ascertained with probabilistic genotyping software (EuroForMix). The motivation for this work was to provide an improved method to generate data to prepare distributions that are used to inform activity level propositions. However, other applications will benefit, particularly those where extraction and quantification are bypassed: For example direct PCR and Rapid DNA technology. The overall aim of this work was to provide a method of quantification that is standardised and can be used to compare results between different laboratories that use different multiplexes. A software solution ”ShinyRFU” is provided to aid calculations.

## 1. Introduction

The notion of using average peak height 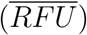 of a DNA profile as a proxy measurement for DNA quantity was introduced by Tvedebrink et al [1, 2], along with a method to apportion quantities per contributor based on analysis of contributor-specific alleles. The method was improved by [3, 4, 5] who used probabilistic genotyping, EuroForMix [6], to determine the relative proportions of known and unknown contributors (*M*_*x*_), that constitute a mixture. The output was used to calculate likelihood ratios given activity level propositions based upon 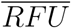. For a review of activity level see Taylor et al. [7]. Here, the authors cite a typical example where the data are discrete and designated as either ‘matching’ or ‘not matching’ a suspect. As with mixture analysis, discrete models (also known as semi-continuous) do not use all of the information available, and have been largely replaced by more efficient continuous models. There is demonstrable consensus that it is preferable to use continuous data and models. However, with respect to activity level, there is no agreement about how such data should be generated. This raises an important question whether results obtained in one laboratory are relevant to another that is using a different protocol? This in turn leads to another obvious question: How can results between different laboratories, using different protocols be compared?

It is not realistic to expect laboratories to agree to use common extraction methods, multiplexes and platforms etc. No two laboratories will be the same; the key is to design a protocol that can be universally applied and facilitates comparisons to be made between laboratories.

Consequently, a methodology has been developed to enable comparisons of DNA profiles generated using different protocols between and within laboratories, so that continuous data can be exchanged and used in Bayesian Networks. This enables activity level propositions to be addressed. The challenge is to provide a standard format that can be easily adopted by all laboratories so that different data-sets may be directly compared and normalised. This is against a background of a variety of different methods, including multiplexes, extraction strategies, equipment etc.

To facilitate, a series of simple calibration control experiments were carried out to establish the relationship between 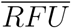 and DNA quantity using regression analysis.

This paper is structured as follows: First there is a description of the current use of qPCR to quantify DNA (section 2), followed by an outline of the important factors that affect the RFU outcome after CE (section 3). The relationship between DNA concentration and average RFU is described in section 4 along with the notion of calibration using control samples. Prior to PCR, samples are diluted to avoid overloading. This affects the outcome and must be compensated with a dilution factor (section 5). To enable comparison between laboratories, it is necessary to normalise data (section 6). There follows a detailed discussion on the limitations of qPCR in section 7. There are general remarks about SWGDAM quality assurance guidelines in relation to applications such as direct PCR and Rapid DNA, where pre-quantification is not possible before PCR (section 8). Finally there is a list of recommendations, and reference to software to assist with calculations (9).

## 2. Use of qPCR to measure DNA quantity

Powerquant^®^ was used to quantify the amount of DNA present in a sample using qPCR [8]. This system is manufactured by Promega. The test is not directed at STRs. Neither the target locus nor its copy number are disclosed by the manufacturer, hence in the following text, the target locus is designated as unknown (*U*). It is advertised as multi-copy. For comparison, an alternative product from Promega is Plexor HY^®^ [9]. This product is disclosed as amplifying a 99bp sequence of a tandemly repeated motif of the RNU2 locus found on chromosome 17. There are 10-20 copies of the repeat sequence per haploid genome [10].

The Powerquant^®^ short amplicon is 81bp (*U*_81_); a longer amplicon of 294bp (*U*_294_) is used to assess levels of degradation. However, it is standard practice to use *U*_81_ to calculate the total amount of DNA present and all quantification values are based upon this marker.

Similar kits are available from different manufacturers such as Quantifiler^®^ Hp and Trio, from Thermo Fisher [11]. These systems detect multi-copy loci with a short amplicon of 80bp and a longer amplicon of 214bp.

## 3. An outline of factors that affect the 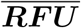 outcome

1. The sampling method: The swabbing or tape technique used to remove cells/ free DNA from the evidence.
2. The extraction method: Technique/ reagents used (solid, magnetic beads etc.)
3. The elution volume (*E*_*V*_): The volume of the eluant in *μ*l that contains the total DNA from the swabs.
4. Quantification: Concentration (*Q*) of DNA in the eluant, using short (*Q*_81_) and long (*Q*_294_) fragments respectively, measured in ng/*μ*l. Variable *Q*_*tot*_ measures the total amount of DNA in *ng* that is recovered from the eluant (*E*_*V*_), hence *Q*_*tot*_ = *Q × E_V_*
5. Degradation factor (*deg*): Measured as the proportion: 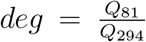 and *deg* ≥ 1 since DNA is subject to varying degrees of degradation, that increasingly affects the high molecular weight loci (see [12], section 4.5.1).
6. The volume of the eluant in *μ*l that is taken for PCR (*T*_*el*_): Laboratories will seek to optimise the amount of DNA that is subject to PCR. A typical target amount may be a total of 1ng. For high template recovery, it is necessary to dilute the sample to obtain the optimum amount, whereas for low template, it may not be possible to achieve the target amount, hence the profile will be lower quality and/or partial.
7. The pre-PCR set-up will consist of:
  a. Volume *T*_*pcr*_: The reaction mix of primers for a given multiplex, *Taq* polymerase and buffer
  b. Volume *T*_*el*_: Taken from the eluant and added to volume of buffer or water (*T*_*dl*_) so that a constant volume *T*_*el_max*_ = *T*_*dl*_ + *T*_*el*_ is always achieved irrespective of the sample analysed. Hence the total PCR volume is *T*_*V*_ = *T*_*pcr*_ + *T*_*dl*_ + *T*_*el*_ and *T*_*V*_ is also a constant volume across all experiments.
8. PCR amplification: The number of cycles; the multiplex used; volumes of buffer; the PCR machine, all affect the outcome.
9. Post-PCR analysis: The analytical platform; manufacturer and model; the injection parameters used; analytical threshold.

## 4. Relationship between DNA concentration and 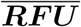

### 4.1. Calibration

The 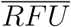 response is dependent upon a number of factors other than the quantity of DNA analysed in the PCR reaction. Different instruments will have different sensitivities. Haas et al [13] showed that there was a big difference between peak heights generated by Genetic Analysers 3500 and 3730 vs. 3130xl. This necessitated that RFUs from the latter were multiplied by a factor of three to standardise the output. Additional dependencies include: the multiplex used; CE injection time, volume, and voltage settings, which must be maintained as constant for the series of experiments or casework using the same conditions. For a given set-up, to be able to convert 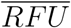 into DNA quantity, it is necessary to carry out calibration using known quantities of undegraded DNA. Furthermore, since there are many variables, this calibration will be laboratory-specific.

### 4.2. Calibration protocol

Fresh blood samples were collected by finger pricks from one donor. Twenty micro-litres of blood was pipetted onto two swabs. The tips of the swabs were cut into an extraction tube. Samples were extracted on the BioRobot EZ1(Qiagen) with the DNA Investigator kit (Qiagen) using the trace protocol and a 200*μ*l elution volume. The samples were quantified with the PowerQuant^®^ System (Promega) and amplified with Promega’s PowerPlex^®^ Fusion 6C System (25 μL reaction volume, 1 ng DNA input, 29 amplification cycles) as recommended by the manufacturer. Amplification was carried out using a Veriti^®^ 96-Well Thermal Cycler (Applied Biosystems™). Samples were injected onto the Applied Biosystems 3500xl Genetic Analyzer at 1.2 kV for 24 s. Results were analysed using the GeneMapper^®^ ID-X Software version 1.6 (Applied Biosystems™)

A dilution series was prepared with input DNA quantities ranging between 0.02 - 22ng in a total of 15*μ*l - the volume used in the PCR reaction (1*μ*l is taken for CE).

DNA concentration (ng/*μ*l) was measured with qPCR using the *U*_81_ amplicon; fluorescence was recorded by the 7500 Fast Real-Time PCR System, Applied Biosystems™. RFU measurements were carried out using conventional capillary electrophoresis (CE) using the 3500xL Genetic Analyzer, Applied Biosystems™. The average RFU value 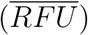 was calculated per DNA profile as:

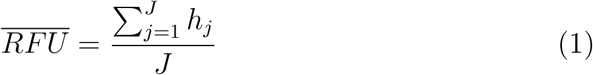

where *h*_*j*_ is the sum of the peak heights per locus *j* = 1,..., *J*, with *J* loci in the multiplex. The formula can further be expanded to include replicates:

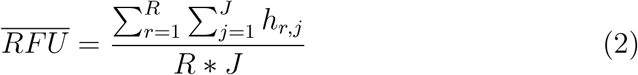

where *r* is the replicate index and *R* is the total number of replicates and replicates are based upon the same multiplex.

The relationship between the two variables is described by a log-linear model (fig 1). A series of concentrations of the undegraded control sample analysed with Fusion 6C [14] ranged between 0.001 - 0.1ng/*μl* in the linear range. As the amount of DNA increases, so does the fluorescent signal. Above 0.1ng/*μl*, the charge-coupled device (CCD) of the CE instrument becomes saturated, and the response is no longer log-linear. A threshold is reached - for Fusion 6C, the non-linear response > 0.1ng/*μ*l resulted in a plateau at 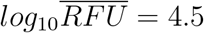.

**Figure 1:**
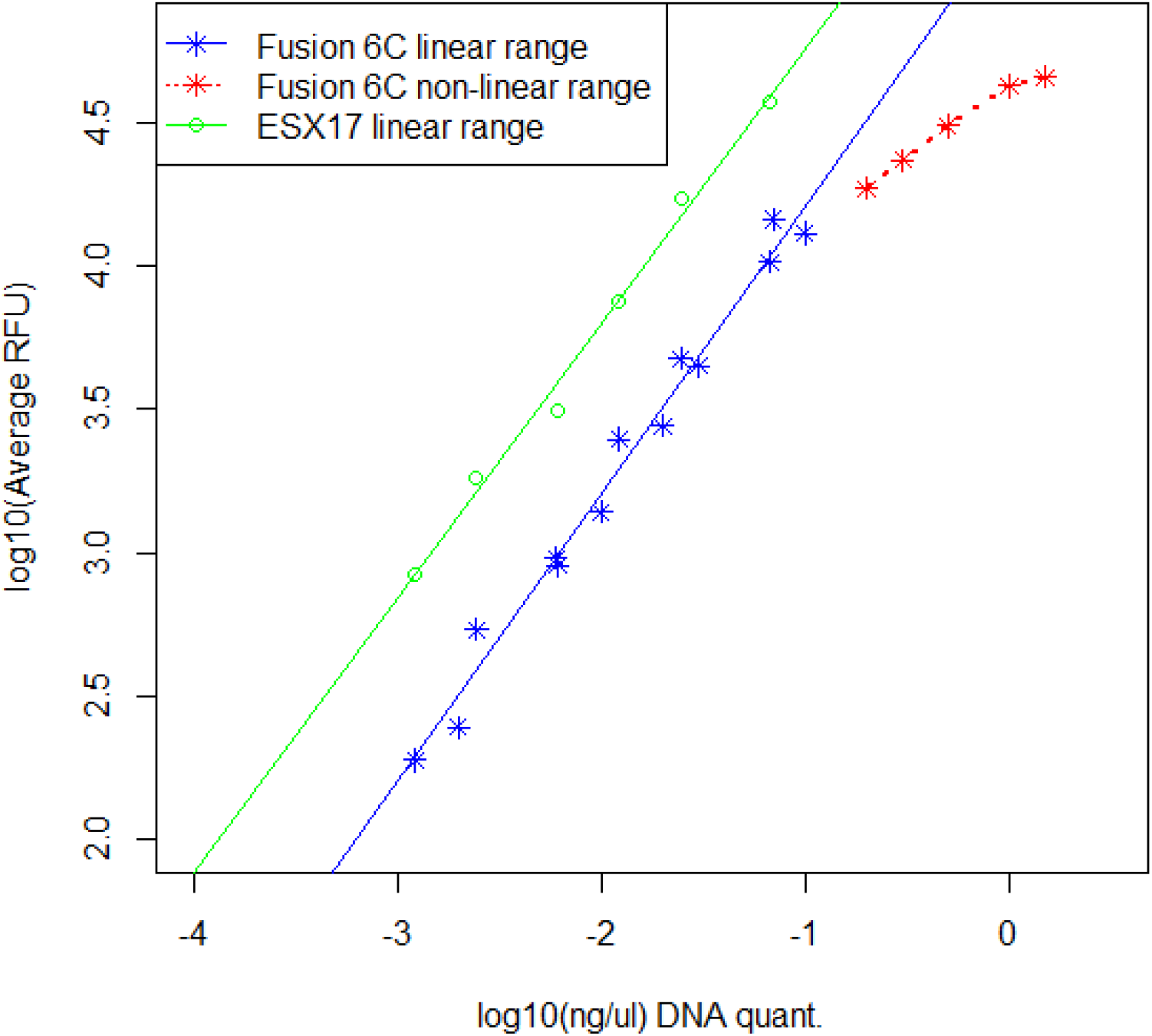
Comparison of *log*_10_(ng/*μ*l) DNA quantification value versus *log*_10_(average RFU) for controls processed using two different multiplexes - Fusion 6C and ESX17

A dilution series using the same control sample analysed with the ESX17 multiplex [15] is shown for comparison. This multiplex elicits a higher fluorescence response for a given DNA quantity compared to Fusion 6C, and is therefore more sensitive. This increased sensitivity is attributed to an additional PCR cycle.

We define an ordinary linear regression model with 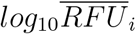 as response and DNA quantity *log*_10_*Q*_*i*_ as explanatory variable, where *i* is an observation:

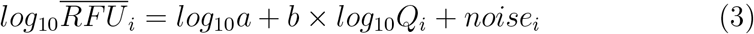

where *noise*_*i*_ is independent identically distributed as normal with expectation zero and constant variance.

The prediction for the response variable for a input DNA quantity *log*_10_*Q*, is then given as the expectation:

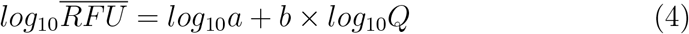

where *a* = intercept and *b* = regression coefficient, which must be estimated. Since the control samples comprise pristine DNA (fig. 1), they provide regressions with very high R-squared values > 0.98. The *b* coefficient is very close to one in both regressions (and standard errors are low). Consequently, by fixing *b* = 1, the prediction equation can be simplified to 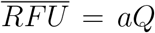; alternatively, to predict the amount of amplifiable DNA for a given 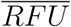, this is simply:

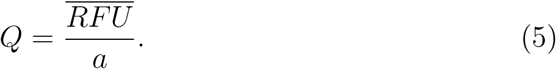

All that is needed to carry out the conversion to DNA quantity is a multiplex specific intercept regression coefficient (*a*) (Table 1).

**Table 1:** Regression analysis i) Fusion 6C and ES17 control samples from fig 1 ii) Vaginal mucosa and skin cells using Fusion 6C analysis. *a* is the intercept and *b* is the regression coefficient. Figures in parentheses for the intercept (*a*) are calculated using fixed *b*=1 (equation: 6). The 95% confidence intervals are provided for the *b* coefficient.

|  | Coefficients |  | Standard Error | 95% CI | R-squared |
| --- | --- | --- | --- | --- | --- |
| Fusion 6C | $\log_{10}a$ | 5.21 (5.2) | 0.08 | | 0.98 |
| | $b$ | 1.00 | 0.04 | [0.91, 1.09] | |
| ESX17 | $\log_{10}a$ | 5.71 (5.8) | 0.09 | | 0.99 |
| | $b$ | 0.96 | 0.04 | [0.84, 1.08] | |
| Vaginal Mucosa | $\log_{10}a$ | 5.08 (5.25) | 0.21 | | 0.57 |
| | $b$ | 0.89 | 0.13 | [0.62, 1.16] | |
| Skin Cells | $\log_{10}a$ | 4.70 (4.84) | 0.20 | | 0.49 |
| | $b$ | 0.91 | 0.13 | [0.66, 1.17] | |

Furthermore, if the regression coefficient is fixed as *b* = 1, then *a* can be estimated by averaging across the *i* = 1,..., *I* observations:

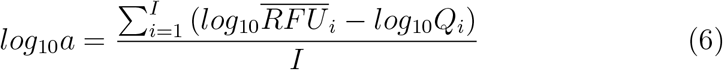

To conclude, provided pristine samples are analysed with minimal degradation, the *U*_81_ amplicon provides an excellent method to estimate the actual amount of amplifiable DNA represented by the multiplex in question and this underpins the basis of the calibration.

#### 4.2.1. Calibration of the regression slope coefficient and intercept

According to Gaigalas and Wang [16], there is an expectation of a perfect log-linear relationship between DNA quantity and fluorescence (measured here as 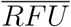), hence *b* = 1. They defined a successful calibration where the observed regression slope fell between 0.95 and 1.05. This condition was met for the tests carried out for Fusion 6C and ESX17 undegraded calibration control samples (fig 1), but not for degraded vaginal mucosa and skin cells (discussed in the next section). The intercept (*a*) coefficient is the theoretical 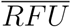 value when 1ng/*μ*l of DNA is tested (it is theoretical since such high quantity would overload the CE instrument such that the RFU response would plateau). It is a measure of the sensitivity of the overall multiplex and CE conditions. Approximately 6 samples are needed for calibration, hence it is easily carried out.

### 4.3. Comparison of casework samples

Data from two separate (Fusion 6C) experiments were compared (Table 1). The first dataset (A) comprised 158 samples of low-level degraded DNA. This dataset was taken from a previous study described in detail by Johannessen et al.[17]. It consists of samples taken in relation to simulated sexual assault: vaginal mucosa and epithelial cells were recovered from fin-gernail swabs, penile swabs and boxer-shorts. The data-set also includes background DNA samples. The second dataset (B) consisted of 118 samples collected from the necks of simulated victim assaults, described in detail by Fantinato et al. [4]: epithelial cells were recovered, that were generally low template and degraded. The results from the two data sets are plotted in fig 2 along with their regressions. Only data from samples that were not diluted were used here.

**Figure 2:**
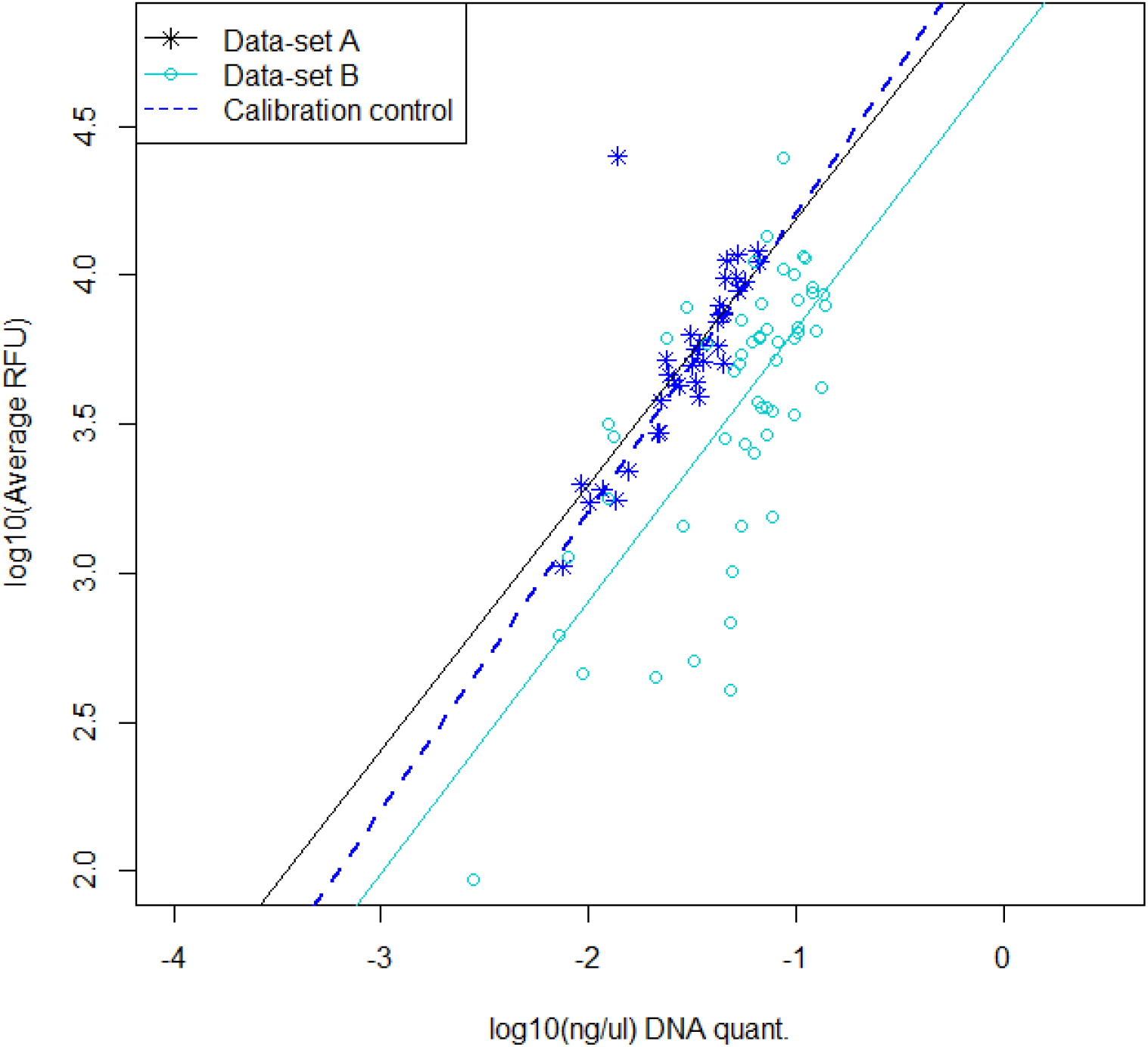
Regressions of data-set A and data-set B processed with Fusion 6C. The control sample calibration regression from fig 1 is superimposed

Apart from a single outlier, for data-set A, a plot of *log*_10_ DNA quantity (*Q* ng/*μ*l) vs. 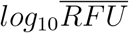 closely followed the control sample regression line from fig. 1. The regression slope (*b* = 0.89) is lower than that achieved by the calibration exercise because degradation causes a spread of samples, increasing the confidence interval which encompasses value *b* = 1

Data-set B also followed the same gradient, but the intercept *a* value was reduced. This is because the DNA was more degraded compared to data-set A (fig. 3); there is a trend for the qPCR quantification test to overestimate the amount of amplifiable DNA, often by an order of magnitude or more.

**Figure 3:**
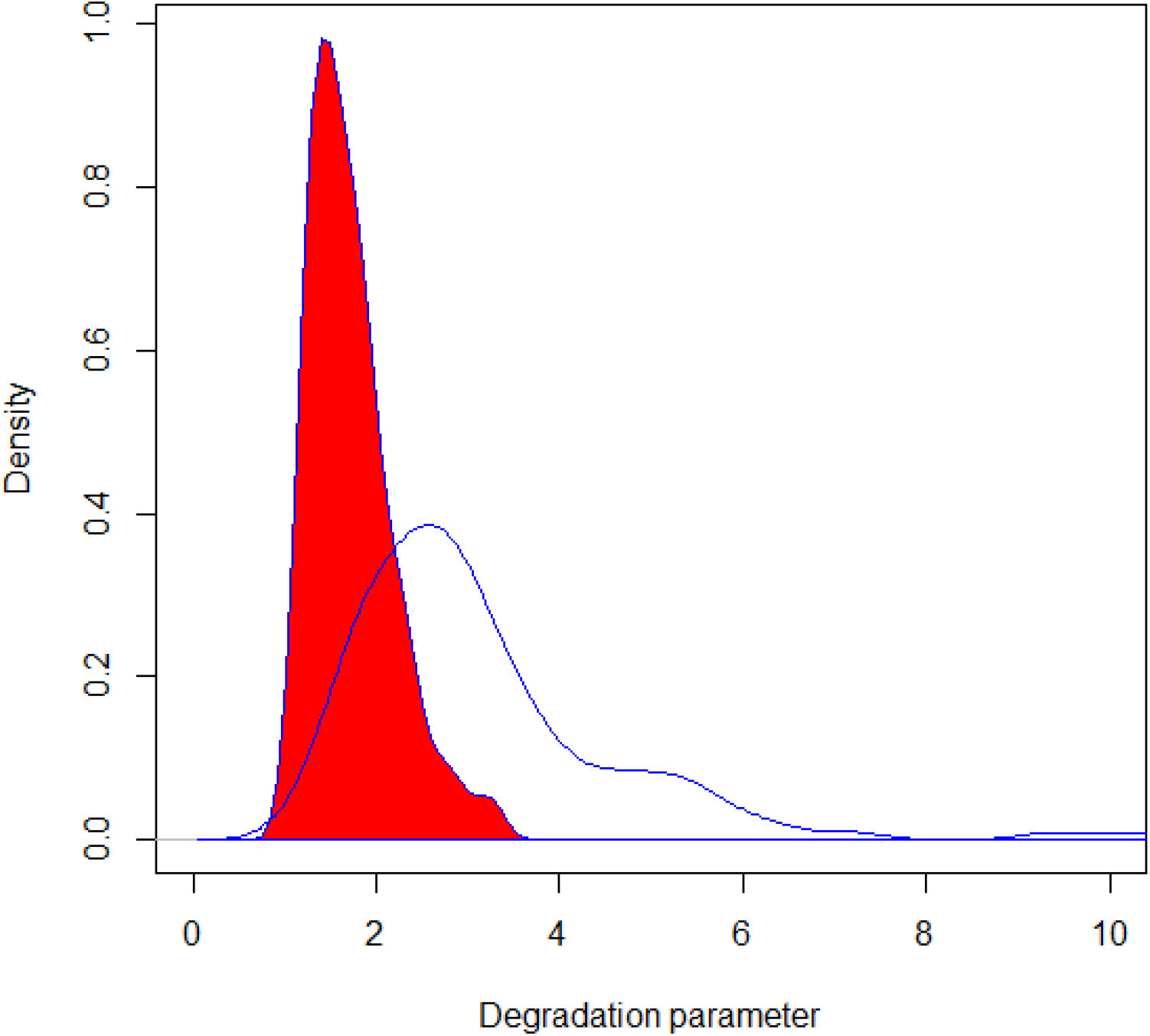
Comparison of the distribution of degradation factors for data-set A (red-filled) and data-set B. Definition of degradation factor (*deg*) is given in section 3

This illustrates the requirement to carry out calibration with undegraded DNA. The intercept coefficient from the calibration study (e.g., *a* = 5.2 for Fusion 6C) is the only parameter required to convert 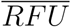 to DNA quantity, regardless of its degraded state (section 6)

## 5. Calculation of the dilution factor

The aim is to provide an optimal amount of DNA into the pre-PCR set-up. If too much DNA is loaded onto a CE instrument, then the signal will be saturated and the profile will be poor quality. This optimum level may vary between laboratories, but is typically 1ng, defined as the total amount of DNA that is provided to the pre-PCR set-up; measured by a quantification method, and provided in a constant volume (*T*_*el_max*_). However, for our subsequent calculations, we must refer to the concentration of DNA that is available in the orginal (undiluted) extract. Furthermore, it is necessary to adjust the calculated 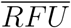 value so that it accurately reflects the expectation from the undiluted stock extract. To achieve this, it is a dilution factor (*dl*) is calculated that is multiplied by the 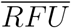 value.

### 5.1. Calculation of the dilution factor when low template DNA is recovered

Refer to section 3 for definitions of the terms used. To make the method clear, examples of calculations are provided in the Supplementary Material (S1.) for the reader to follow. In order to provide the optimum quantity for the PCR reaction, *T*_*el*_ is taken from elution volume and added to *T*_*dl*_ of dilution buffer or water. The dilution factor is calculated:

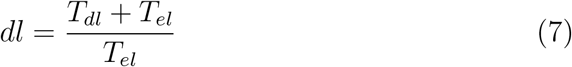

If *T*_*el*_ = *T*_*el_max*_ then *dl* = 1

### 5.2. Calculation of the dilution factor when high template DNA is recovered

If the DNA sample is high template, then it may be necessary to carry out a double dilution. Volume *T*_*el*_ from the eluant is added to volume *T*_*dl*2_ of dilution buffer/water. For the second dilution, volume *V*_*el*_ is taken and added to the PCR set-up volume of *T*_*dl*_. The dilution factor is calculated:

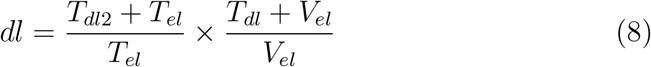

A worked example is provided in the Supplementary Material (S1.2.)

Fig 4 shows results of both data-sets when the dilution factor (*dl*) is applied. The Fusion 6C control calibration regression line is superimposed. Data-set A has much higher quantities compared to data-set B, hence dilution factors applied were much higher. The adjusted data also follow the control sample calibration regression line. Note that the high 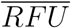 calculated values > 10^4^, exceed the saturation threshold in the CE instrument and would not be physically observed. The effect of applying the dilution factor is to preserve the log-linearity of the response.

**Figure 4:**
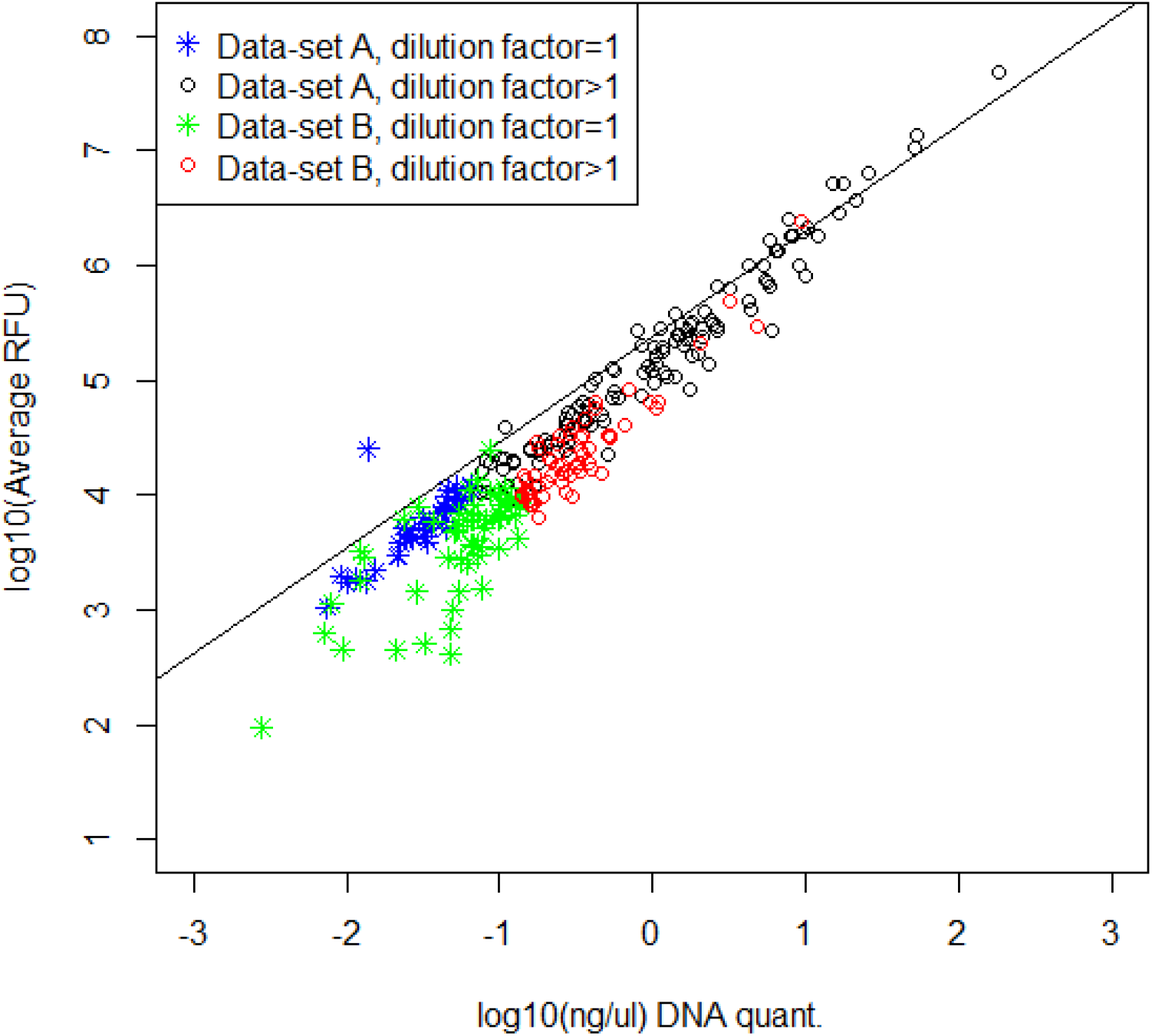
Comparison of *log*_10_*Q* (ng/*μ*l) DNA quantification value versus 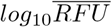 for data-sets A and B analysed with Fusion 6C. The data are identified according to whether the dilution factor *dl* = 1 or *dl* > 1 (see section 5). The Fusion 6C control calibration regression line is superimposed

The calculated 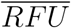 is multiplied by the dilution factor to give the adjusted value. If the sample is a mixture, then it is also multiplied by the mixture proportion of the person of interest (POI) as described by [3] to give:

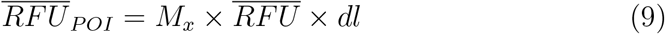

## 6. Normalisation of data to compare results generated from different protocols

In order for laboratories to adopt data from other laboratories, it is necessary to ensure that necessary adjustments are made. For example, the ESX17 multiplex is more sensitive than Fusion 6C, hence if a series of experiments is carried out using the latter, a lab employing Fusion 6C may be interested to use such data to evaluate results given activity level propositions. For a given DNA quantity, ESX17 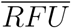 results will be greater than those from Fusion 6C, hence will not be directly comparable. In order to facilitate comparisons of data from different laboratories that use a variety of different methods and procedures, e.g., multiplexes, it is necessary to carry out normalisation. To do this, a common standard is needed, the Fusion 6C data shown in Fig 1 are adopted for this demonstration. The ESX17 are normalised to the Fusion 6C data as follows: The predicted concentration of DNA (*Q*) for given 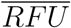 results, based on the regression equation where the coefficients have been pre-determined (section 4), is given as:

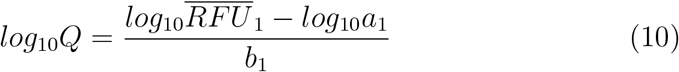

where 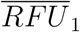, *a*_1_ and *b*_1_ are the Fusion 6C observation and pre-determined coefficients; 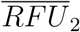, *a*_2_ and *b*_2_ are the ESX17 observation and pre-determined coefficients. To normalise ESX17 to the Fusion 6C expectation, for any given value of *Q*:

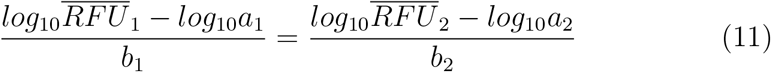

By rearrangement, 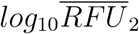 is normalised:

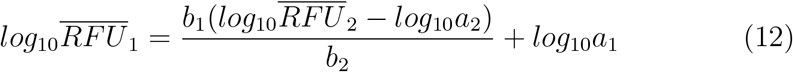

In this example, both *b*_1_ and *b*_2_=1, hence the equation simplifies to:

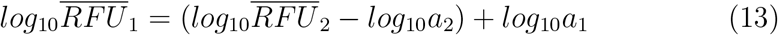

or without logs:

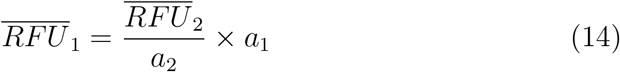

Finally, it is necessary to take into account the different volumes that may be extracted from swabs in order to provide the elution volume (*E*_*V*_). Samples used to create data-set A were extracted into a volume of *E*_*V*_ = 100*μl*, whereas those used to create data-set B were extracted into *E*_*V*_ = 200*μl*, consequently the latter is diluted 1:2.

It follows that if the total quantities (ng) of DNA extracted were identical in an experiment, the concentration for data-set B will be half that of the data-set A. Proceeding to remove identical volumes *T*_*el*_ into the PCR set-up will result in data-set B giving a 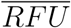 value that is half that of data-set A. To compensate, the *E*_*V*_ extraction volume is normalised to be the same for all samples, so that DNA concentrations are based upon an adjusted (standardised) volume of *E*_*V*_ =100*μ*l. If 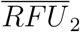 is the value for data-set B experiment, the normalised (adjusted) value 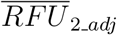 is calculated:

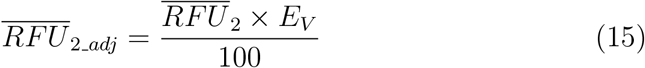

i.e., 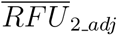 substitutes 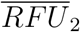 in equation 14.

The same calculation is carried out to normalise the DNA concentration:

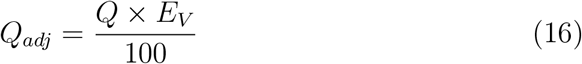

Because *E*_*V*_ = 100*μl* is used as the standard volume, everything is normalised against this (figs. 2, 4 and 6 all show adjusted values).

**Figure 5:**
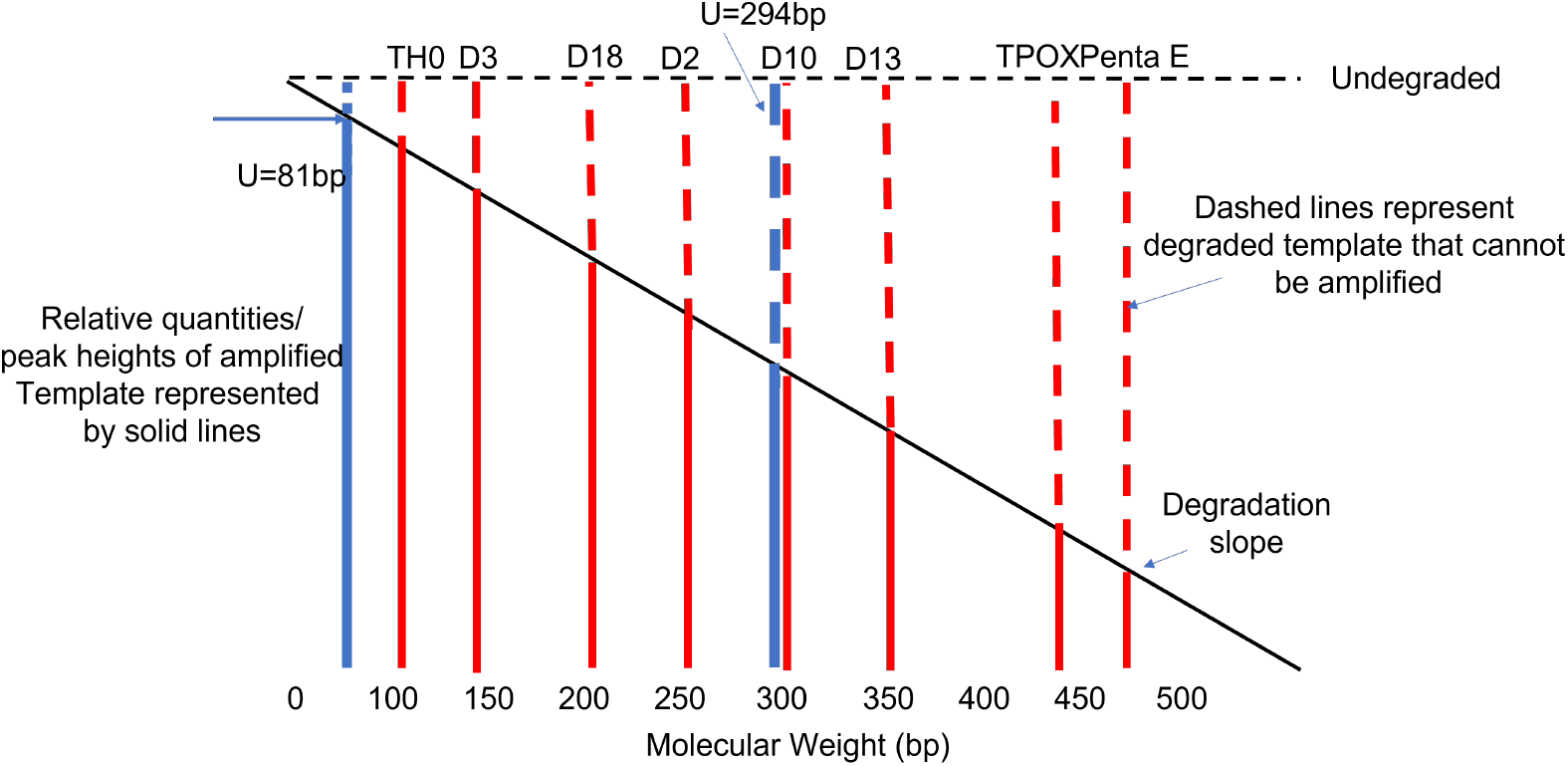
Schematic illustration: there are a small selection of n loci shown relative to their approximate molecular weights in the Fusion 6C multiplex. The height of the solid bars give the relative quantity recorded per locus in a degraded sample.The *U*_81_ locus over-estimates quantities of all loci, whereas the *U*_294_ locus over-estimates quantities at D13, TPOX and PentaE and under-estimates TH0, D3, D18 and D2.

**Figure 6:**
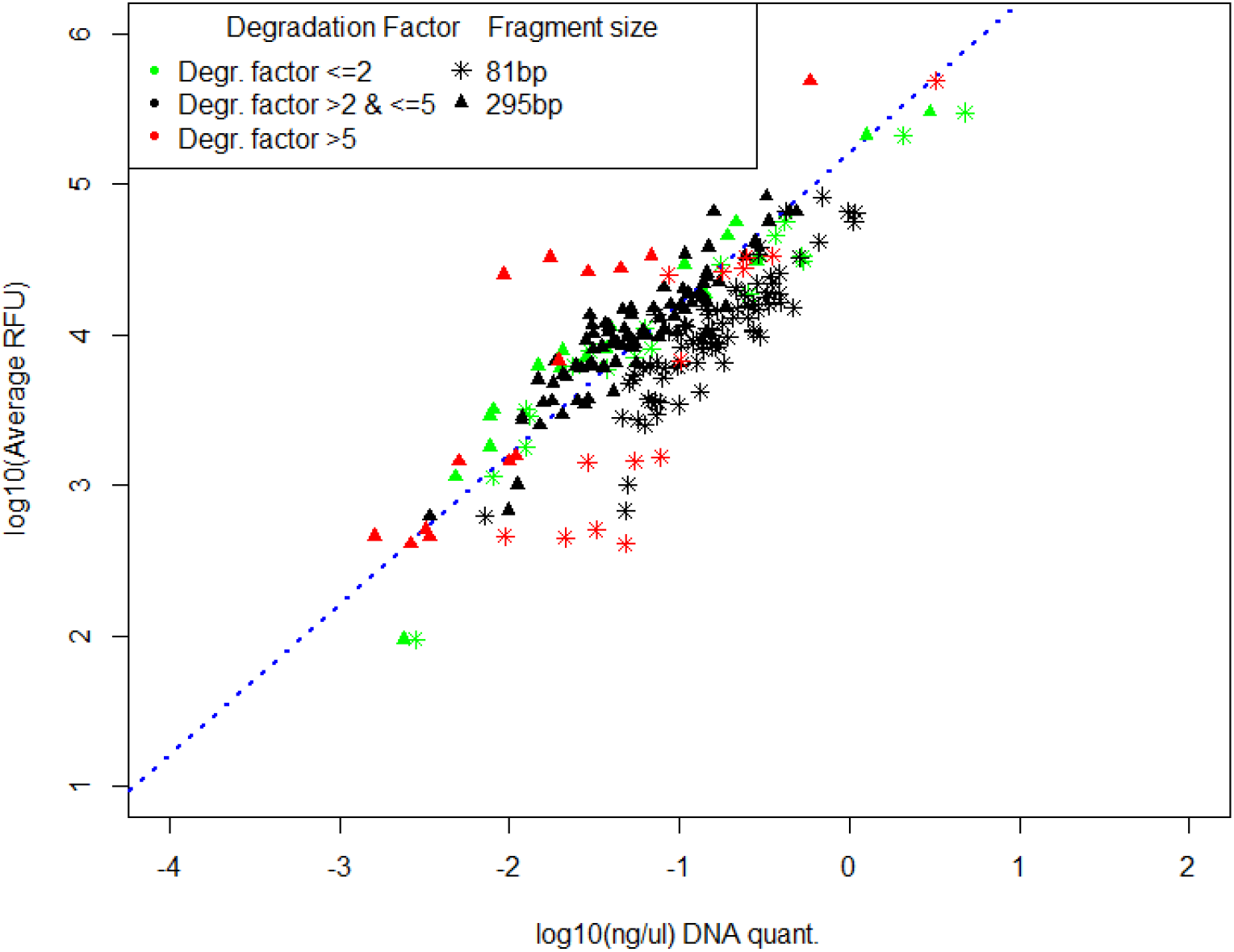
Effect of degradation factor on quantification estimates based on the short 81bp fragment and the long 295bp fragment. The control calibration regression line is shown

### 6.1. Using total quantity

However, a preferable alternative, is to instead calculate the total amount in ng of DNA (*Q*_*tot*_) recovered in the elution volume. From equation 10, with *b* = 1:

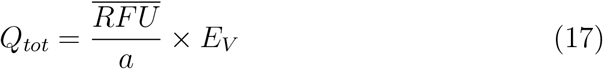

Both approaches described are equivalent to each other, but the *Q*_*tot*_ method has advantages in that it is simpler to calculate; results are given in absolute ng rather than ng/*μ*l, and normalisation is not necessary. Consequently it should be amenable to standardisation. Example calculations are provided in the Supplementary Material, showing that conversions between the methods described are easily accomplished.

### 6.2. Extraction efficiency

In addition to the elution volume, the method of extraction e.g. fluid, solid or magnetic beads is expected to have an impact as DNA recoveries differ between protocols. Wang et al. [18] compared five different DNA/RNA extraction protocols, showing differences could be quantified.

We have not included an extraction efficiency factor here, but further work to describe a protocol (eq. 17) is planned: For example, known volumes of blood placed upon swabs and dried, and subsequently subjected to the laboratory’s standard extraction method would enable regression parameters (Fig 1) that translated the peak heights all the way back to the amount of DNA on the sampled item. Consequently, even the extraction method of a study would not need to align with a laboratory in order to make use of the data. This idea is similar to that proposed by Taylor et al [19], where distributions for sampling and extraction efficiency were developed and used to back-calculate DNA quantity on the original object.

## 7. Limitations of qPCR quantification

With respect to the Powerplex Fusion 6C kit, with 23 autosomal loci, alleles (*A*_1..*k*_) fall within a range of 70-470bp [14]. The Powerplex *U*_81_ short amplicon will amplify all fragmented locus *U* DNA that is > 81bp, provided that the fragmentation has not taken place within the amplicon itself. With undegraded DNA, the quantity of DNA (*Q*_81_) equals the estimated actual value of the averaged multiplexed fragment peak heights 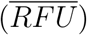 (equation 5). Consequently, *Q* = *Q*_81_ where *Q* is the observed quantity from the multiplex fragments. However, when degradation has occurred, this affects the probability that a target fragment will amplify (fig. 5). As the molecular weight of the target increases, then the probability of amplification decreases. The *U*_81_ fragment amplifies all *U* fragments *A* > 81bp; it does not compensate for the reduction in quantification values for high molecular weight of *A*_1..*k*_ multiplexed markers. Consequently, *Q*_81_ > *Q*. Conversely, the high molecular *U*_295_ marker is approximately mid-way in the Fusion 6C range. It cannot amplify fragments *U* < 295bp, hence this will result in an underestimate of the amplifiable quantity of *A*_1..*k*_ < 295*bp* DNA present. On the other hand, since all fragments *U* > 295bp will be amplified, this results in over-estimation of *Q* in the range *A*_1..*k*_ = 295 − 500bp, where amplification rates decrease with increasing molecular weight.

This leads to the consequence, that for moderately degraded DNA, the under- and over-estimations of the *U*_294_ fragment tend to balance each other (fig. 6), leading to an estimate of DNA quantity that is closer to *Q*, whereas for highly degraded DNA where the majority of fragments are *A*_1..*k*_ < 295bp, there will be substantial underestimation of *Q*.

Whereas the Powerquant method simply produces a global value based on the quantity of fragments of a single locus *U*_81_, the peak height quantification method takes account of the relative amounts of allele specific molecular weights *A*_1..*k*_ of the Fusion 6C and other multiplexes.

Fig 6 shows the effect of basing the quantification value on *U*_81_ vs *U*_295_ amplicons relative to the degradation factor (*deg*). The calibration regression line *log*_10_*Q* vs. 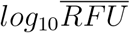 from the Fusion 6C control samples (Fig. 1) is overlaid as a dashed line. For degraded material, the points for the *U*_81_ amplicon are displaced to the right of the calibration regression line, showing overestimated values (up to an order of magnitude). When measurements are made relative to the *U*_295_ amplicon, points are shifted to the left and are much closer to the calibration regression line. This will happen when the average MW of markers ≈ 295bp. Shifts are more extreme for high levels of degradation (*deg* > 5), placing points to the left of the calibration regression line.

In order to help overcome the ‘noise’ in the relationship between quantification and average peak height, an option would be to take degradation into account using expected values of *h*_*j*_ from the exponential model suggested by Taylor et al. [20]; then model the regression.

## 8. General applicability where prior quantification is precluded

This focus of this study was to propose a method to standardise a data collection method applicable to activity level propositions. However, the method can be used wherever there is a need to carry out quantification of a sample. Standard 9.4 of the SWGDAM Quality Assurance Standards for Forensic DNA Testing Laboratories [21]: *”the laboratory shall quantify or otherwise calculate the amount of human DNA in forensic samples prior to nuclear DNA amplification”*. Accordingly, many laboratories have standard operating procedures that stipulate a lower threshold quantity that must be reached in order to report test results. However, for reasons outlined in section 7, using standard qPCR methods, a *prior* assessment of the total *amplifiable* amount of DNA present will likely be underestimated, especially when the DNA is degraded. Quantification is best carried out following nuclear DNA amplification and CE. Some methods: direct PCR and Rapid DNA technology by-pass the extraction and quantification methods [22]. Consequently, quantification can only be carried out *a posteriori* using eq. 17. Since there is no elution volume, it is necessary to prepare wet or dry control samples, where a known quantity of DNA is applied to a swab (or other material). If the protocol requires a wet swab, then controls are applied in a given volume of fluid; otherwise for a dry swab, they should be dried: preparation of controls must always emulate the working protocol. For the calibration exercise, controls are applied directly to the PCR set-up. The calibration plot is in absolute ng rather than ng/*μ*l. It is also unnecessary to use probabilistic genotyping to assign mixture proportions if only an overall value is required.

## 9. A summary of recommendations

1. Any given system can be calibrated with a dilution series of approximately six control samples that must be undegraded and of known quantity. They must be carefully selected to be within the log-linear range of fluorescence response as illustrated by fig 1. A successful calibration will give regression slope *b* = 1 (±0.05, allowance for error). The intercept (*log*_10_*a*) coefficient is a measurement of the sensitivity of the method.
2. The amount of sample amplified is adjusted so that an optimum amount is loaded to the PCR. To calculate the concentration of DNA in the original elution, a dilution factor is calculated as described in section 5, eqs. (7) and (8)
3. The 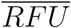 ng/*μ*l value is apportioned between contributors by multiplying by the mixture proportion (*M*_*x*_) and the dilution factor (*dl*). *M*_*x*_ is calculated using probabilistic genotyping software
4. The total quantity (*Q*_*tot*_) in ng of DNA recovered is calculated with eq. 17
5. Worked examples are shown in the Supplementary Material S1

## 10. Software to calculate average RFU

Version 3.4.1 of EuroForMix http://www.euroformix.com/ will automatically calculate 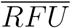. In addition, a user friendly Shiny application https://shiny.rstudio.com/ has been prepared. The program utilises EuroForMix, but automates the analysis of multiple samples placed in a single file. The program, along with a user manual and example data is available at https://github.com/peterdgill/ShinyRFU.

## 11. Conclusion

The aim of this study was to provide a method to standardise data collection across different protocols that are practised within and between laboratories. The relationship between DNA quantity and 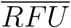 is demonstrated. Fixing the regression slope coefficient (*b* = 1), justified by calibration study, simplifies the method to calculate predicted values of 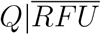 based on the predetermined value of the *a* coefficient (equation 5). There are many papers published that describe observations that are used to inform probabilities given activity level propositions. Different laboratories use different protocols, so the question is whether data generated by a laboratory can be used by another that uses different protocols (equipment such as genetic analyser, multiplexes, extraction methods etc.). It is clearly not possible to standardise protocols between laboratories, but it is possible to compare results provided that the protocols are characterised. It was demonstrated how this can be achieved by using the regression intercept (*a*) coefficient, extraction volumes, and volumes taken for PCR. There is room for further improvement, particularly by taking extraction efficiency into consideration.

Finally, it is clear that qPCR using kits such as Powerquant^®^ are useful prior indicators of DNA quantity and quality, allowing an assessment of the extraction volume to be forwarded to PCR. However, because qPCR does not provide a direct measurement of the relative quantity of DNA of the multiplex utilised, it will tend to underestimate the levels of amplifiable degraded DNA - typified by data-set B. Consequently the average RFU 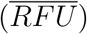 method, to estimate the total quantity of DNA (*Q*_*tot*_) recovered, is suggested as a way forward to standardise results between different protocols run by different laboratories.

## Acknowledgements

The authors are grateful to Helen Johannessen for supplying dataset A and to Chiara Fantinato for supplying dataset B.

## Supplementary material

### S1. Dilution Factor Calculations

#### S1.1. Example 1

In this example, a stain is extracted into a total elution volume *E*_*V*_ = 100*μ*l. A portion is taken for quantification using qPCR and the concentration is recorded as *Q*_*i*_ = 0.01ng/*μ*l or a total of *Q*_*tot*_ = 0.01 *×* 100 = 0.5ng. This value serves as a guide to optimise the amount of DNA forwarded to the PCR set-up. With this example, 1ng total is optimal.

In the PCR set-up, the total PCR volume *T*_*V*_ = 50*ul* where *T*_*pcr*_ = 35*ul* consists of PCR mastermix and primers. The remainder of *T*_*dl*_ + *T*_*el*_ = 15*ul* consists of water and DNA template respectively. Since the qPCR estimate is 0.01ng/*μ*l, it is only possible to take a maximum of 15*μ*l x 0.01ng/*μ*l = 0.15ng in total i.e. no water is added to the PCR set up volume with this example.

It is assumed that all PCR amplifications are carried out using equal total PCR volume *T*_*V*_ = *T*_*pcr*_ + *T*_*el*_ + *T*_*dl*_. Further more, the CE injection parameters are assumed to be constant.

##### S1.1.1. Calculation of the dilution factor

The dilution factor (*dl*) is calculated:

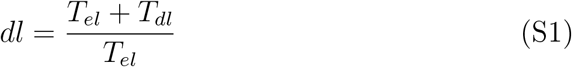

Following the example:

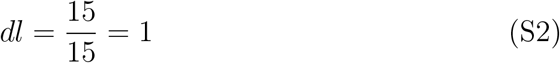

##### S1.1.2. Example 2

Taking the same variables as for Example 2, if the recovery of DNA is *Q*_*i*_ = 0.3*ng/μ*l, to avoid overloading the PCR reaction, it is necessary to take a dilution of *T*_*el*_ = 3*μ*l: *T*_*dl*_ = 12*μ*l in order to achieve the optimum 1ng template DNA. Hence, from equation S1:

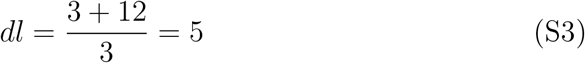

#### S1.2. Example 3

With this example, there is a large amount of DNA recovered where *Q*_*i*_ = 2ng/*μ*l. The optimum 1ng is therefore contained in just 0.5*μ*l which is difficult to accurately aliquot. Therefore a portion of the eluant is diluted twice in order attain the desired 1ng template for the PCR set-up. The dilution factor is calculated:

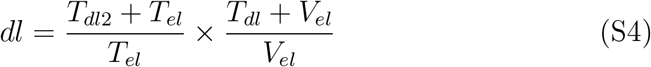

For the first dilution, an aliquot of *T*_*el*_ = 1*μ*l is diluted by the addition of *T*_*dl*2_ = 9*μ*l water,resulting in an estimated 0.2 ng/*μ*l of template. In the second round of dilution we take 1ng template which is in *V*_*el*_ = 5*μ*l, added to *T*_*dl*_ = 10*μ*l so that a total of 15*μ*l is added to the PCR set-up.

The dilution factor is calculated from eq. S4:

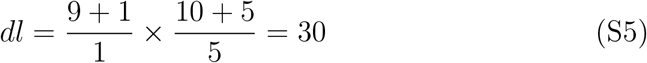

### S2. Calculation of the DNA quantity

To calculate the 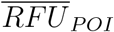 adjusted by the mixture proportion (*M*_*x*_), and dilution factor (*dl*):

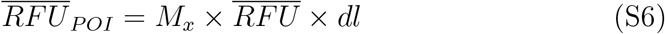

Then the total quantity of DNA recovered is calculated by dividing 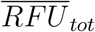 by the regression intercept of the multiplex used, multiplied by the elution volume:

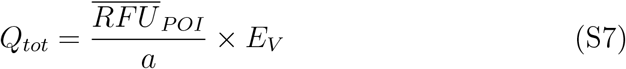

To calculate the total DNA quantity, the examples all assume Fusion 6C multiplex, *a* = 5.21, and *M*_*xPOI*_ = 0.5, and an observed 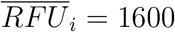

#### S2.1. Example 1

Here *dl* = 1. The POI adjusted value from eq. S6 is:

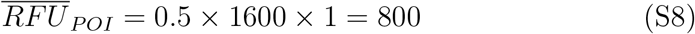

The total quantity of DNA recovered from the POI in the elution volume *E*_*V*_ = 100*ul* from eq. S7 is:

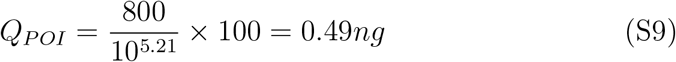

#### S2.2. Example 2

Here *dl* = 5. The POI adjusted value from eq. S6 is:

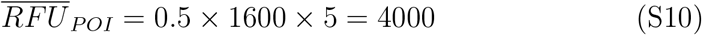

The total quantity of DNA recovered from the POI in the elution volume *E*_*V*_ = 100*ul* from eq. S7 is:

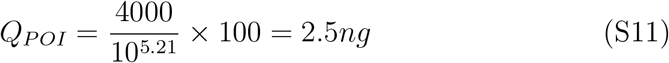

#### S2.3. Example 3

Here *dl* = 30. From eq. S6

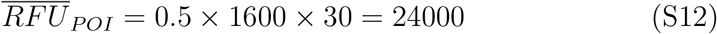

and the total quantity of DNA recovered from the POI in the elution volume from eq. S7 is:

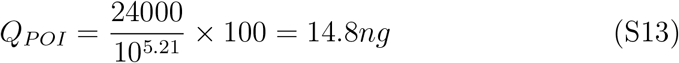

