## Supplementary Material for "Limitations of qPCR to estimate DNA quantity: An RFU method to facilitate inter-laboratory comparisons for activity level, and general applicability"

### S1. Dilution Factor Calculations

#### *S1.1. Example 1*

In this example, a stain is extracted into a total elution volume $E_V = 100\mu$l. A portion is taken for quantification using qPCR and the concentration is recorded as $Q_i = 0.01$ng/$\mu$l or a total of $Q_{tot} = 0.01 \times 100 = 0.5$ng. This value serves as a guide to optimise the amount of DNA forwarded to the PCR set-up. With this example, 1ng total is optimal.

In the PCR set-up, the total PCR volume $T_V = 50ul$ where $T_{pcr} = 35ul$ consists of PCR mastermix and primers. The remainder of $T_{dl} + T_{el} = 15ul$ consists of water and DNA template respectively. Since the qPCR estimate is 0.01ng/$\mu$l, it is only possible to take a maximum of 15$\mu$l x 0.01ng/$\mu$l = 0.15ng in total i.e. no water is added to the PCR set up volume with this example.

It is assumed that all PCR amplifications are carried out using equal total PCR volume $T_V = T_{pcr} + T_{el} + T_{dl}$. Further more, the CE injection parameters are assumed to be constant.

##### *S1.1.1. Calculation of the dilution factor*

The dilution factor ($dl$) is calculated:

$$dl = \frac{T_{el} + T_{dl}}{T_{el}} \tag{S1}$$

Following the example:

$$dl = \frac{15}{15} = 1 \tag{S2}$$

##### *S1.1.2. Example 2*

Taking the same variables as for Example 2, if the recovery of DNA is $Q_i = 0.3ng/\mu$l, to avoid overloading the PCR reaction, it is necessary to take a dilution of $T_{el} = 3\mu$l : $T_{dl} = 12\mu$l in order to achieve the optimum 1ng template DNA. Hence, from equation S1:

$$dl = \frac{3 + 12}{3} = 5 \tag{S3}$$

*S1.2. Example 3*

With this example, there is a large amount of DNA recovered where $Q_i = 2\text{ng}/\mu\text{l}$. The optimum 1ng is therefore contained in just $0.5\mu$l which is difficult to accurately aliquot. Therefore a portion of the eluant is diluted twice in order attain the desired 1ng template for the PCR set-up. The dilution factor is calculated:

$$dl = \frac{T_{dl2} + T_{el}}{T_{el}} \times \frac{T_{dl} + V_{el}}{V_{el}} \tag{S4}$$

For the first dilution, an aliquot of $T_{el} = 1\mu$l is diluted by the addition of $T_{dl2} = 9\mu$l water,resulting in an estimated 0.2 ng/$\mu$l of template. In the second round of dilution we take 1ng template which is in $V_{el} = 5\mu$l, added to $T_{dl} = 10\mu$l so that a total of $15\mu$l is added to the PCR set-up.

The dilution factor is calculated from eq. S4:

$$dl = \frac{9+1}{1} \times \frac{10+5}{5} = 30 \tag{S5}$$

### S2. Calculation of the DNA quantity

To calculate the $\overline{RFU}_{POI}$ adjusted by the mixture proportion ($M_x$), and dilution factor ($dl$) :

$$\overline{RFU}_{POI} = M_x \times \overline{RFU} \times dl \tag{S6}$$

Then the total quantity of DNA recovered is calculated by dividing $\overline{RFU}_{tot}$ by the regression intercept of the multiplex used, multiplied by the elution volume:

$$Q_{tot} = \frac{\overline{RFU}_{POI}}{a} \times E_V \tag{S7}$$

To calculate the total DNA quantity, the examples all assume Fusion 6C multiplex, $a = 5.21$, and $M_{xPOI} = 0.5$, and an observed $\overline{RFU}_i = 1600$

*S2.1. Example 1*

Here $dl = 1$. The POI adjusted value from eq. S6 is:

$$\overline{RFU}_{POI} = 0.5 \times 1600 \times 1 = 800 \tag{S8}$$

The total quantity of DNA recovered from the POI in the elution volume $E_V = 100ul$ from eq. S7 is:

$$Q_{POI} = \frac{800}{10^{5.21}} \times 100 = 0.49ng \tag{S9}$$

*S2.2. Example 2*

Here $dl = 5$. The POI adjusted value from eq. S6 is:

$$\overline{RFU}_{POI} = 0.5 \times 1600 \times 5 = 4000 \tag{S10}$$

The total quantity of DNA recovered from the POI in the elution volume $E_V = 100ul$ from eq. S7 is:

$$Q_{POI} = \frac{4000}{10^{5.21}} \times 100 = 2.5ng \tag{S11}$$

*S2.3. Example 3*

Here $dl = 30$. From eq. S6

$$\overline{RFU}_{POI} = 0.5 \times 1600 \times 30 = 24000 \tag{S12}$$

and the total quantity of DNA recovered from the POI in the elution volume from eq. S7 is:

$$Q_{POI} = \frac{24000}{10^{5.21}} \times 100 = 14.8ng \tag{S13}$$
